# Human fibroblasts from aged individuals exhibit chromosomal instability through replication stress caused by oxidative stress

**DOI:** 10.1101/2025.06.19.660647

**Authors:** Kailin Zhu, Guan Chen, Yueyi Ren, Kenji Iemura, Kozo Tanaka

**Affiliations:** Department of Molecular Oncology, Institute of Development, Aging and Cancer (IDAC), Tohoku University, 4-1 Seiryo-machi, Aoba-ku, Sendai, Miyagi 980-8575, Japan

**Keywords:** Aging, Chromosomal instability, Human fibroblasts, Oxidative stress, Replication stress

## Abstract

Aneuploid cells are known to increase with age. Previously, we demonstrated that aneuploid cells increase in fibroblasts from aged mice due to chromosomal instability (CIN), which is caused by oxidative stress. It is unclear whether this phenomenon also occurs in human cells, which are more resistant to oxidative stress than mouse cells. Here, we found that fibroblasts from aged individuals exhibited an increase in aneuploid cells. The frequency of chromosome missegregation and micronuclei increased in these cells, indicating CIN. A DNA fiber assay revealed the presence of replication stress, accompanied by an increase in 53BP1 nuclear bodies and ultrafine bridges. Increased levels of reactive oxygen species derived from mitochondria, along with reduced mitochondrial membrane potential, imply that these cells experienced oxidative stress due to mitochondrial functional decline. Antioxidant treatment reduced the frequency of chromosome missegregation and micronuclei, suggesting that oxidative stress causes CIN. Oxidative stress also causes replication stress, which precedes CIN. Spindle microtubules were stabilized in fibroblasts from aged individuals, which was alleviated by antioxidant treatment. Taken together, these findings suggest that aging-related CIN in human fibroblasts is caused by oxidative stress associated with mitochondrial dysfunction, which induces replication stress that in turn causes CIN through microtubule stabilization. Although human fibroblasts are more resistant to the ambient oxygen environment than mouse fibroblasts, our findings showed that they undergo oxidative stress that causes CIN with age in a manner similar to mouse fibroblasts, revealing a conserved phenomenon in mammalian cells.

## INTRODUCTION

Genomic instability, a key characteristic of aging, manifests as a gradual build-up of genetic damage over an organism’s lifespan. This damage encompasses various forms, such as point mutations, translocations, gains and losses of chromosomes, telomere shortening, and gene disruption (Lopez-Otin et al., 2013). Among them, chromosomal gains and losses, known as aneuploidy, is known to increase with age (Ricke & van Deursen, 2013). While chromosome aberration in oocytes associated with maternal age is a well-known age-related chromosome abnormality (Nagaoka et al., 2012), an increase in aneuploid cells in somatic cells with aging has also been reported (Macedo et al., 2017). For example, it is known that Y chromosome is frequently lost in males with increasing age (Jacobs et al., 1963; Pierre & Hoagland, 1972). In the brain, mosaic aneuploidy increases with age and has been associated with neurodegenerative diseases (Andriani et al., 2017). With increasing age, mice also exhibited an increase in chromosome abnormalities, including aneuploidy (Lushnikova et al., 2011). Cancer is also characterized by genomic instability (Hanahan & Weinberg, 2011), and aneuploidy is observed in most cancer cells (Weaver & Cleveland, 2006). Chromosomal instability (CIN), a state of frequent chromosome missegregation, typically accompanies aneuploidy in cancer cells (Bakhoum & Compton, 2012; Gordon et al., 2012; Tanaka & Hirota, 2016). Recently, we reported that primary fibroblasts isolated from aged mice exhibit aneuploidy and CIN (Chen et al., 2023). Fibroblasts from aged mice showed elevated levels of reactive oxygen species, along with a decline in mitochondrial function, suggesting oxidative stress. The rates of chromosome missegregation in cells from aged mice was reduced by treatment with antioxidants. We identified replication stress as a cause of CIN in aged mouse cells, which may promote CIN through microtubule stabilization. Our findings revealed the age-related onset of CIN and indicated a novel link between oxidative stress and CIN during aging. However, it is not known whether the same phenomenon occurs in human cells. While primary human fibroblasts proliferate for 50 or more population doublings under atmospheric oxygen environment, primary mouse fibroblasts rapidly enter a period of declining growth rate (Parrinello et al., 2003), suggesting that mouse cells are more susceptible to oxidative stress than human cells (Hornsby, 2003). In contrast to human cells that stop dividing with a normal karyotype in most of the cases (Lansdorp, 2000), mouse cells undergo immortalization with chromosomal abnormalities after the period of slowed growth (Todaro & Green, 1963), raising a question on the assumption that age-related aneuploidy occurs via a common pathway in mouse and human cells.

To study aneuploidy and CIN with age in human cells, we observed fibroblasts isolated from young and aged individuals. We found that fibroblasts isolated from aged individuals exhibit aneuploidy and CIN. Similar to fibroblasts from aged mice, fibroblasts from aged individuals are under oxidative stress, which causes CIN through replication stress and microtubule stabilization. Our data show that the link between oxidative stress and age-related CIN is conserved between mouse and human cells despite the significant difference in their sensitivity to oxidative stress.

## RESULTS

### Human fibroblasts isolated from aged individuals exhibit CIN

To study how chromosomal stability changes with age in human cells, we obtained human fibroblasts from the NIA Aging Cell Repository at the Coriell Institute for Medical Research. We selected four cell lines derived from young individuals and four from aged individuals (Table 1). Of these, two cell lines from young individuals and four cell lines from aged individuals were skin fibroblasts. We also selected two lung fibroblasts from fetal tissues to see the difference depending on the cellular origins. We first observed the proliferation of these cell lines. As shown in Fig. 1A, the proliferation rate of fibroblasts from aged individuals was slower than that of fibroblast from young individuals. In subsequent experiments, we used fibroblasts at a population doubling level (PDL) ranging from 17 to 35, during which the cells grew at a constant rate. We did not use cells at PDL 36 or higher, where cell growth was markedly suppressed. We then examined the chromosome numbers of these cell lines in metaphase spreads. We found a significant increase in metaphases with abnormal chromosome numbers in fibroblasts from aged individuals (Fig. 1B), in line with the notion that aneuploid cells increase with age. Both the number of metaphases with fewer and more than 46 chromosomes increased, while the average number of chromosomes remained unchanged (Fig. S1A, B), showing that despite the marked increase in aneuploid cells from aged individuals, the modal number did not change. To study whether CIN exists in fibroblasts from aged individuals as a cause of aneuploidy, we first observed the proportion of interphase cells with micronuclei, which are formed as a result of chromosome missegregation, in fixed cell samples. We found that fibroblasts from aged individuals showed a higher rate of the appearance of micronuclei (Fig. 1C). We next quantified the percentage of chromosome missegregation in fixed cell samples. Fibroblasts from aged individuals showed higher rates of lagging chromosomes and chromosome bridges than those from young individuals (Fig. 1D). These data suggest that fibroblasts from aged individuals exhibit CIN. We found that skin and lung fibroblasts from young individuals showed comparable levels of micronucleation and chromosome missegregation rates. These cells exhibited a similar phenotype in all subsequent experiments, so they were analyzed together. To further study CIN in fibroblasts from aged individuals, we performed live cell imaging and observed chromosome missegregation during mitosis. We confirmed that chromosome missegregation, both lagging chromosomes and chromosome bridges, were increased in fibroblasts from aged individuals (Fig. 1E, F). We found that the total duration of prometaphase (from nuclear envelope breakdown (NEBD) to chromosome alignment) as well as the duration of mitosis (from NEBD to nuclear decondensation) were increased in fibroblasts from aged individuals (Fig. 1G). To see whether the activity of the spindle assembly checkpoint (SAC), which delays anaphase onset until all the chromosomes attach to spindle microtubules, differ in fibroblasts from young and aged individuals, we compared the duration of mitosis in the presence of nocodazole, a microtubule destabilizer, or Eg5 inhibitor III, which inhibits the activity of a motor protein Eg5, both of which arrest cells in mitosis by activating the SAC. As shown in Fig. S1C, the duration of mitotic arrest was comparable between fibroblasts from young and aged individuals both in the presence of nocodazole and Eg5 inhibitor III, showing that the SAC activity is not altered in fibroblasts from aged individuals. We also observed the cellular response after prolonged mitotic arrest. When cells are arrested in mitosis for a prolonged period, they either die in mitosis (mitotic cell death) or enter the next cell cycle (mitotic slippage). It is known that whether cells undergo mitotic cell death or mitotic slippage after prolonged mitotic arrest varies not only between cell lines, but also within a population of the same cell line (Gascoigne & Taylor, 2008). We found that the rate of mitotic slippage did not differ between fibroblasts from young and aged individuals in the presence of nocodazole as well as in the presence of Eg5 inhibitor III (Fig. S1D).

**Fig. 1.**
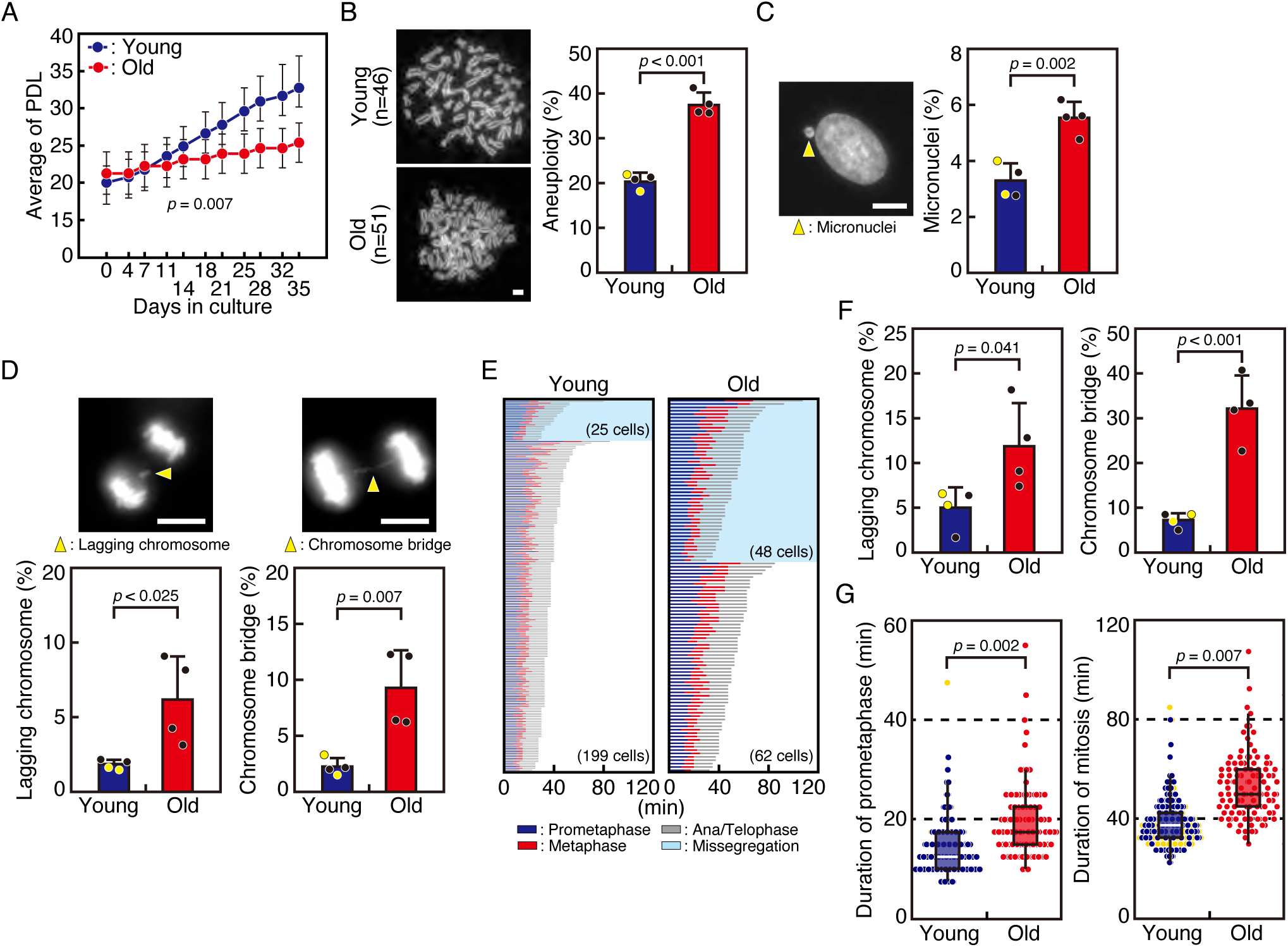
Human fibroblasts isolated from aged individuals exhibit CIN. (A) Growth of human fibroblasts isolated from young and aged individuals. Fibroblasts from young and aged individuals (n = 4 each) were seeded at 1 × 10^6^ cells/10 cm dish and cultured for 3-7 days. At confluence, the cells were trypsinized and counted by TC20 automated cell counter (Bio-Rad). Population doubling level (PDL) was calculated using the formula PDL = 3.33 × [log (number of cells harvested) – log (number of cells seeded)] + (doubling level at seeding). Error bars represent S.D. Repeated-measures ANOVA was used to judge the significance. (B) Chromosome numbers of fibroblasts isolated from young and aged individuals. 50-105 metaphase chromosome spreads of fibroblasts from young and aged individuals (n = 4 each) were observed. Representative images of metaphase chromosome spreads of cells from young and aged individuals are shown in the left panels. Scale bar: 5 μm. Percentages of aneuploid cells were shown in the right graph. Black-filled circles represent skin fibroblasts, while yellow-filled circles represent lung fibroblasts. Error bars represent S.D. P values were obtained using the Student’s *t*-test. (C) Micronucleation rates of fibroblasts isolated from young and aged individuals. Rates of fibroblasts with micronuclei from young and aged individuals (n = 4 each) were fixed and stained with DAPI. Scale bar: 5 μm. 504-631 cells per cell line were analyzed. A representative image of a micronucleus is shown in the left panel (arrowhead). Black-filled circles represent skin fibroblasts, while yellow-filled circles represent lung fibroblasts. Error bars represent S.D. P-value was obtained using the Welch’s *t*-test. (D) Chromosome missegregation rates of fibroblasts isolated from young and aged individuals. Fibroblasts from young and aged individuals (n = 4 each) were treated as in (C). Representative images of a lagging chromosome and a chromosome bridge are shown in the upper panels (arrowheads). Rates of lagging chromosomes and chromosome bridges were quantified for 46-66 cells per cell line. Scale bars: 5 μm. Black-filled circles represent skin fibroblasts, while yellow-filled circles represent lung fibroblasts. Error bars represent S.D. P values were obtained using the Student’s *t*-test. (E) Mitotic progression of fibroblasts isolated from young and aged individuals. Fibroblasts from young and aged individuals (n = 4 each, 22-61 cells per cell line) were treated with SiR-DNA and subjected to live cell imaging. Images were taken every 2.5 min for 96 h, and mitotic progression of individual cells was tracked and displayed in different colors depending on the mitotic phases. Cells that underwent chromosome missegregation are separately shown, and number of cells in each category is indicated. (F) Chromosome missegregation rates of fibroblasts isolated from young and aged individuals. Rates of lagging chromosomes and chromosome bridges in fibroblasts from young and aged individuals (n = 4 each) shown in (E) are plotted. Black-filled circles represent skin fibroblasts, while yellow-filled circles represent lung fibroblasts. Error bars represent S.D. P-values were obtained using the Student’s *t*-test. (G) Mitotic duration of fibroblasts isolated from young and aged individuals. Time from nuclear envelope breakdown to chromosome alignment (duration of prometaphase) and time from nuclear envelope breakdown to cytokinesis (duration of mitosis) in fibroblasts from young and aged individuals (n = 4 each, 22-61 cells per cell line) shown in (E) are shown as box and dot plots. Red- and blue-filled circles represent skin fibroblasts, while yellow-filled circles represent lung fibroblasts. Error bars represent S.D. P-values were obtained using the Mann–Whitney *U* test.

**Table 1.**
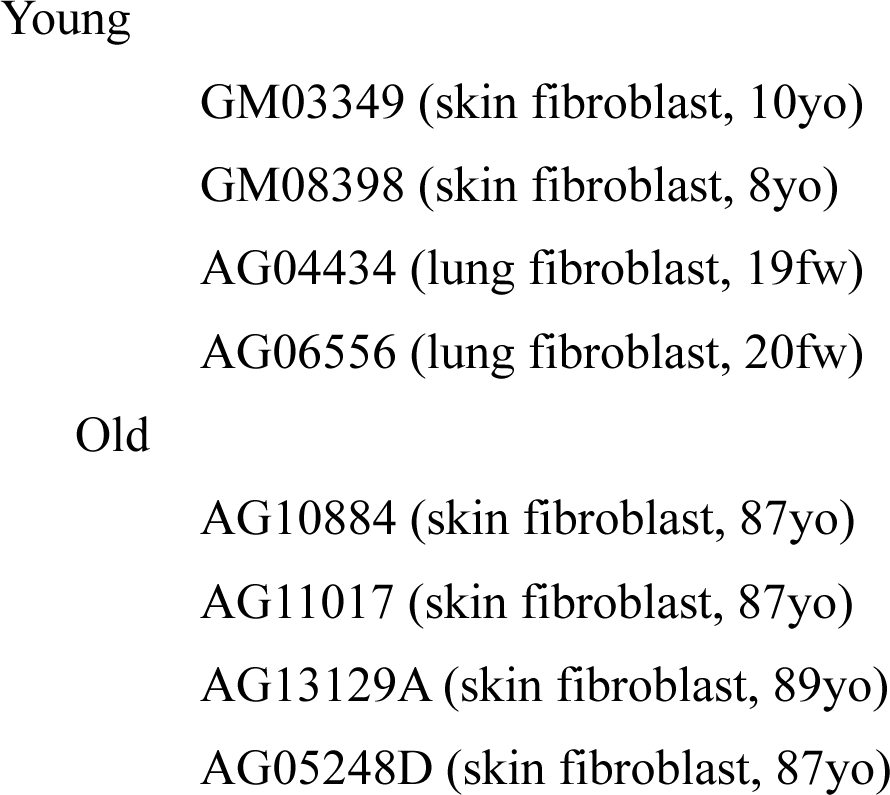
Human fibroblast cell lines used in this study.

Since senescent cells are known to increase with age (McHugh & Gil, 2018), we examined their percentage using SPiDER-βGal, a fluorescent marker of senescence (Doura et al., 2016). The percentage of cells positive for the SPiDER-βGal signal increased in fibroblasts from aged individuals, as expected. (Fig. S1E). However, the presence of micronuclei and the SPiDER-βGal signal were not significantly correlated in cells from both young and aged individuals (Fig. S1E), implying that micronucleation and cellular senescence are independent phenomena. It is also known that myofibroblasts, which are differentiated from fibroblasts to facilitate tissue repair, increase in lung fibroblasts from aged individuals (Paxson et al., 2011; Plikus et al., 2021). Quantifying the percentage of cells positive for α-smooth muscle actin (α-SMA), a myofibroblast marker, revealed an increase in fibroblasts from aged individuals (Fig. S1F). Examining the relationship between micronucleation and differentiation into myofibroblasts revealed no correlation in fibroblasts from both young and aged individuals (Fig. S1F), suggesting that micronucleation is unrelated to differentiation to myofibroblasts.

### Human fibroblasts isolated from aged individuals are under replication stress

Next, we observed the presence of DNA damage in fibroblasts from young and aged individuals. The number of γ-H2AX foci, which represents the presence of DNA double strand breaks (DSBs) (Mah et al., 2010), was increased in fibroblasts from aged individuals (Fig. 2A), consistent with previous reports (C. Wang et al., 2009). We then observed cells with 53BP1 foci, which represent DSBs under processing (Mirman & de Lange, 2020), and found that they were increased in fibroblasts from aged individuals (Fig. 2B). We noticed that these 53BP1 foci were large and few in number, suggesting that they are 53BP1 nuclear bodies formed around DNA lesions generated by replication stress (Lukas et al., 2011). It was reported that 53BP1 nuclear bodies are formed in G1 phase as a result of replication stress in the previous cell cycle. Consistent with the idea that fibroblasts from aged individuals are under replication stress, the percentage of cells with ultrafine bridges (UFBs), which are thin DNA threads connecting sister chromatids in anaphase and can result from replication stress (Baumann et al., 2007; Chan et al., 2007; Chan et al., 2009), was increased in fibroblasts from aged individuals (Fig. 2C). Furthermore, metaphase chromosome spreads showed an increase in chromosome breaks, which are indicative of structural chromosome aberrations that arise from pre-mitotic defects including replication stress, in fibroblasts from aged individuals (Fig. 2D). To directly assess DNA replication, we performed a DNA fiber assay, in which the progression of individual replication forks and the asymmetry between sister replication forks were measured (Halliwell et al., 2020). Fibroblasts from aged individuals showed significantly slower fork rates than those from young individuals (Fig. 2E). Furthermore, asymmetric sister fork progression was increased in fibroblasts from aged individuals (Fig. 2F), indicating impaired replication fork progression. Collectively, these data suggest that fibroblasts from aged individuals are under replication stress. As we previously reported that fibroblasts from aged mice are also under replication stress (Chen et al., 2023), replication stress may commonly increase with age in mammalian cells.

**Fig. 2.**
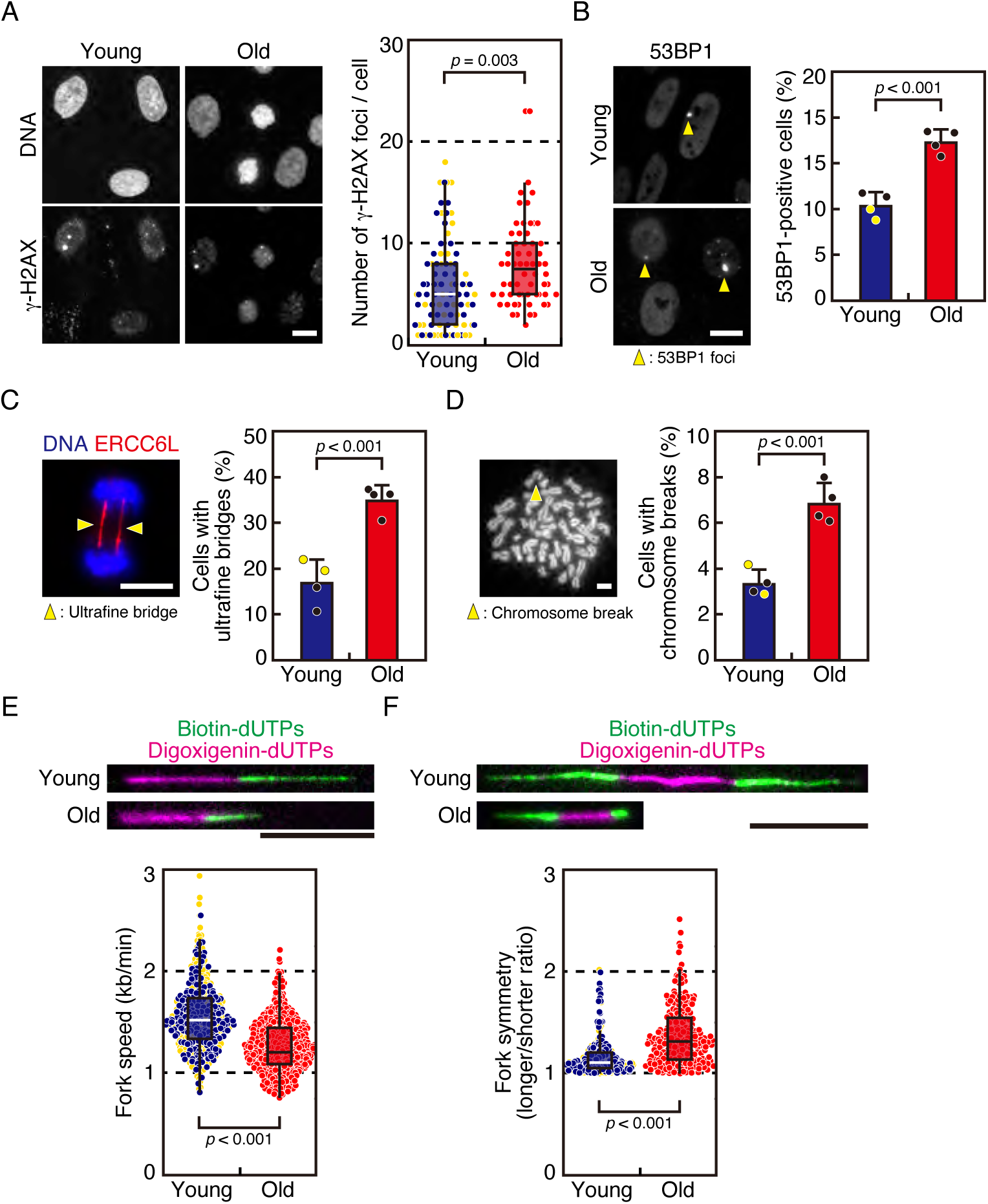
Human fibroblasts isolated from aged individuals are under replication stress. (A) Number of γ-H2AX foci in fibroblasts isolated from young and aged individuals. Fibroblasts from young and aged individuals (n = 4 each) were fixed and stained with an antibody against γ-H2AX. DNA was stained with DAPI. Representative images are shown. Scale bar: 5 μm. Numbers of γ-H2AX foci were quantified for 15-24 cells per cell line and displayed as box and dot plots in the right graph. Data of cells from different cell lines are shown in different colors. Red- and blue-filled circles represent skin fibroblasts, while yellow-filled circles represent lung fibroblasts. Error bars represent S.D. P values were obtained using the Mann-Whitney *U*-test. (B) Rates of cells with 53BP1 nuclear bodies in fibroblasts isolated from young and aged individuals. Fibroblasts from young and aged individuals (n = 4 each) were fixed and stained with an antibody against 53BP1. Representative images of 53BP1 nuclear bodies (arrowheads) were shown in the left panels. Scale bar: 5 μm. Rate of cells with 53BP1 nuclear bodies were quantified for 95-138 cells per cell line. Black-filled circles represent skin fibroblasts, while yellow-filled circles represent lung fibroblasts. Error bars represent S.D. P-value was obtained using the Welch’s *t*-test. (C) Rates of cells with ultrafine bridges in fibroblasts isolated from young and aged individuals. Fibroblasts from young and aged individuals (n = 4 each) were treated with RO-3306 for 12 h, released and fixed after 1 h, and stained with an antibody against PICH (ERCC6L). DNA was stained with DAPI. A representative image of ultrafine bridges (arrowheads) is shown in the left panel. Scale bar: 5 μm. 46-66 cells were counted per cell line. Error bars in the right graph represent S.D. Black-filled circles represent skin fibroblasts, while yellow-filled circles represent lung fibroblasts. P-value was obtained using the Student’s *t*-test. (D) Chromosome breaks in fibroblasts isolated from young and aged individuals. 50-105 metaphase chromosome spreads of fibroblasts from young and aged individuals (n = 4 each) were observed. A representative image of a chromosome break (arrowhead) is shown in the left panel. Scale bar: 5 μm. Percentages of cells with chromosome breaks were shown in the right graph. Error bars represent S.D. Black-filled circles represent skin fibroblasts, while yellow-filled circles represent lung fibroblasts. P values were obtained using the Student’s *t*-test. (E) Fork speed in fibroblasts isolated from young and aged individuals. Fibroblasts from young and aged individuals (n = 4 each) were sequentially pulse-labeled with 200 μM digoxigenin-conjugated deoxyuridine and 200 μM biotin-conjugated deoxyuridine for 10 min each. DNA fiber spreads were prepared from the labeled cell suspension, and stained with Anti-Digoxigenin-Rhodamine, Fab fragments and Streptavidin, Alexa Fluor^®^ 488 Conjugate. Representative fibers are shown. Scale bar: 5 μm. Replication fork speeds were determined through the length of both labels (digoxigenin-dUTPs + biotin-dUTPs) for 101-138 fibers per cell line, according to the previous report that 1 μm is approximately equal to 3.5 kb of DNA (Luebben et al., 2014). 101-138 fibers per cell line were quantified and shown as box and dot blots. Error bars represent S.D. Red- and blue-filled circles represent skin fibroblasts, while yellow-filled circles represent lung fibroblasts. P-value was obtained using the Mann–Whitney *U* test. (E) Fork symmetry in fibroblasts isolated from young and aged individuals. Cells were treated as in (D). Representative fibers are shown. Scale bar: 5 μm. Fork symmetry was determined by dividing longer biotin-dUTPs label on a bi-directional replication fork with shorter biotin-dUTPs label for 49-79 fibers per cell line and shown as box and dot blots. Error bars represent S.D. Red- and blue-filled circles represent skin fibroblasts, while yellow-filled circles represent lung fibroblasts. P-value was obtained using the Mann– Whitney *U* test.

### Fibroblasts from aged mice are under oxidative stress

In a previous study, we reported that fibroblasts from aged mice are under oxidative stress, which promotes CIN (Chen et al., 2023). To determine whether human fibroblasts from aged individuals are also under oxidative stress, we examined the level of reactive oxygen species (ROS) in human fibroblasts by detecting the CellRox Green signal. We found that the ROS level was higher in fibroblasts from aged individuals (Fig. 3A). The specificity of the signal was confirmed by its reduction when cells were treated with N-acetylcysteine (NAC), an antioxidant (Aruoma et al., 1989; Burgunder et al., 1989) (Fig. 3A). As a major origin of ROS production, we observed the level of superoxide in mitochondria using MitoSOX Red and found that it was higher in fibroblasts from aged individuals (Fig. 3B). The MitoSOX Red signal was reduced when cells were treated with NAC, as expected. The reduction of CellROX and MitoSOX signals by NAC treatment even in cells from young individuals indicates that they are under oxidative stress in an atmospheric oxygen environment. It is known that increased ROS production is caused by mitochondrial functional decline, which is related to aging (Passos et al., 2010; Vigneron & Vousden, 2010). We then investigated mitochondrial function by analyzing the mitochondrial membrane potential by comparing the fluorescence signal ratios of aggregated versus monomeric JC-10. As shown in Fig. 3C, the ratio of aggregated to monomeric JC-10 was lower in fibroblasts from aged individuals, indicating that mitochondrial membrane potential was reduced. These data suggest that mitochondrial functional decline results in increased ROS production in fibroblasts from aged individuals, consistent with our previous results in mouse primary fibroblasts (Chen et al., 2023). Interestingly, NAC treatment increased the mitochondrial membrane potential (Fig. 3C), possibly due to the protective effect of NAC on mitochondrial function, as reported previously (Zheng et al., 2024; Zhou et al., 2021).

**Fig. 3.**
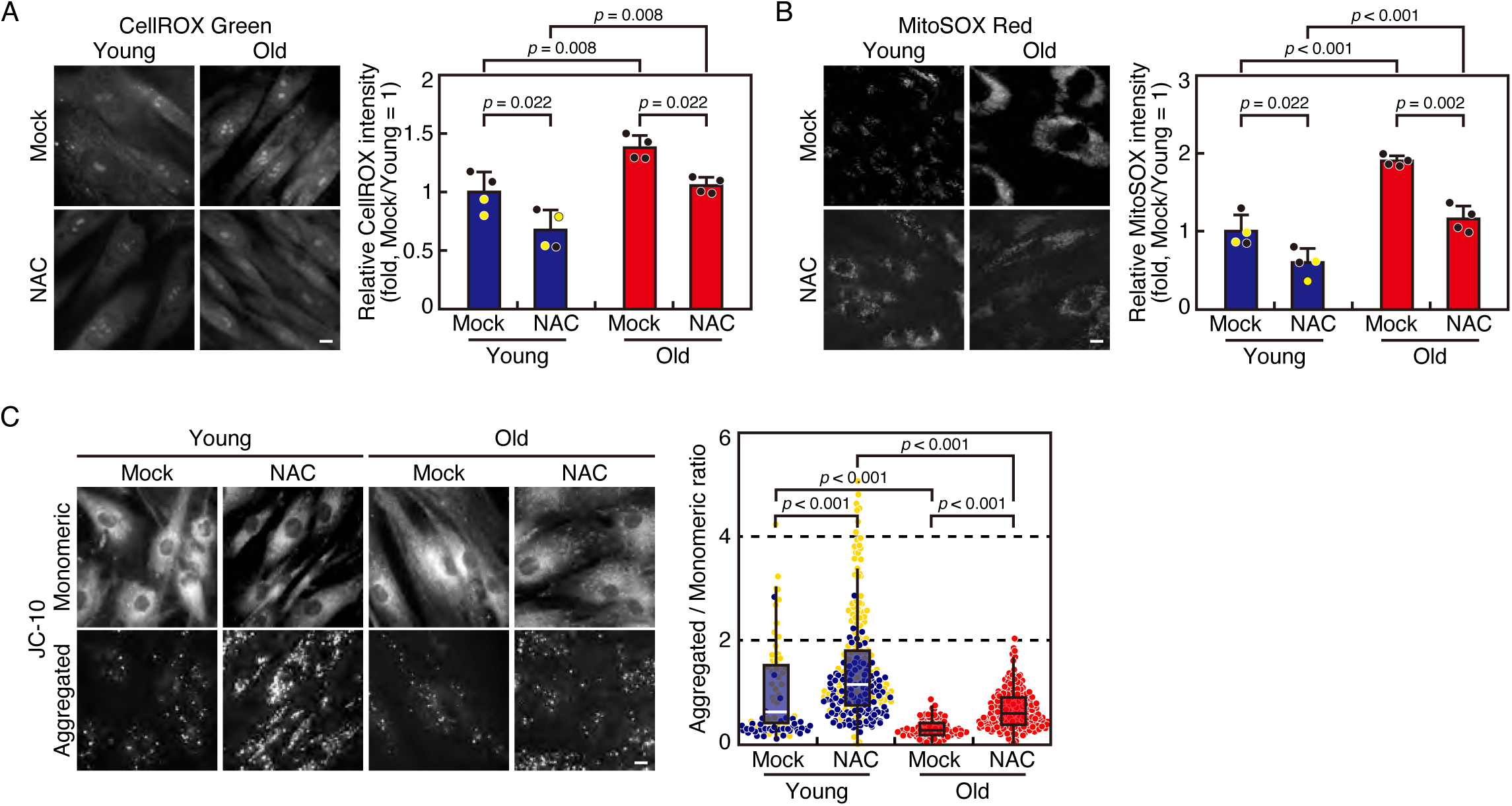
Human fibroblasts from aged individuals are under oxidative stress. (A) Detection of ROS in fibroblasts isolated from young and aged individuals in the presence or absence of an antioxidant. Fibroblasts from young and aged individuals (n = 4 each) were treated with or without NAC for 48 h. Then cells were incubated with CellROX green reagent for 30 min before fixation and observed under microscope. Representative images were shown in the left panels. Scale bar: 5 μm. Quantification of the ROS level is shown in the right graph. Fluorescence intensity of 17-35 cells per cell line were quantified. The average intensity of cells from young individuals without NAC was set as 1. Black-filled circles represent skin fibroblasts, while yellow-filled circles represent lung fibroblasts. Error bars represent S.D. P values were obtained using the Tukey–Kramer multiple comparison test. (B) Detection of superoxide in fibroblasts isolated from young and aged individuals in the presence or absence of an antioxidant. Fibroblasts from young and aged individuals (n = 4 each) were cultured and treated as in (A). Then cells were incubated with MitoSOX Red reagent for 10 min and observed under microscope. Representative images were shown in the left panels. Scale bar: 5 μm. Quantification of the superoxide level is shown in the right graph. Fluorescence intensity of 27-60 cells per cell line was quantified. The average intensity of mock-treated cells from young individuals was set as 1. Black-filled circles represent skin fibroblasts, while yellow-filled circles represent lung fibroblasts. Error bars represent S.D. P values were obtained using the Tukey–Kramer multiple comparison test. (C) Detection of mitochondrial membrane potential in fibroblasts isolated from young and aged individuals in the presence or absence of an antioxidant. Fibroblasts from young and aged individuals (n = 4 each) were cultured and treated as in (A). Then cells were incubated with JC-10 reagent for 30 min and observed under microscope. The signal representing JC-10 monomeric form (emission at 520 nm), and the signal representing JC-10 aggregated form upon polarization of mitochondrial membrane (emission at 590 nm) are shown. Representative images are shown in the left panels. Scale bar: 5 μm. Quantification of the level of mitochondrial membrane potential, shown as the ratio of the emission at 590 nm to that at 520 nm, is shown in the right graph. The values of 33-98 cells per cell line were quantified and displayed as box and dot plots. The average ratio of mock-treated cells from young individuals was set as 1. Red- and blue-filled circles represent skin fibroblasts, while yellow-filled circles represent lung fibroblasts. Error bars represent S.D. P value was obtained using the Steel-Dwass multiple comparison test.

### Antioxidant treatment ameliorates CIN in human fibroblasts

To examine the causal relationship between oxidative stress and CIN in fibroblasts from aged individuals, we observed micronucleation rates in the presence or absence of NAC. As shown in Fig. 4A, the percentage of cells with micronuclei was reduced in the presence of NAC in fibroblasts from both young and aged individuals, although it was not statistically significant in the former, showing that increased oxidative stress is involved in the increase in micronucleation rates. We also quantified the percentage of chromosome missegregation in fixed cell samples in the presence of NAC. We found that the rate of chromosome missegregation was significantly reduced by NAC treatment in fibroblasts from both young and aged individuals (Fig. 4B), further demonstrating that oxidative stress is a cause of CIN in human fibroblasts. The observation that the chromosome missegregation rate was reduced by NAC treatment even in fibroblasts from young individuals suggests that the atmospheric oxygen environment causes oxidative stress that leads to chromosome missegregation. To verify this possibility, we observed chromosome missegregation in cells cultured under a 3% oxygen concentration. We found that the chromosome missegregation rates were lower in fibroblasts from both young and old individuals then in those cultured under an atmospheric oxygen concentration (Fig, 4C), indicating that the atmospheric oxygen environment causes CIN through oxidative stress. When cells under 3% oxygen were treated with NAC, the chromosome missegregation rate was reduced in fibroblasts from aged individuals, but not in fibroblasts from young individuals (Fig. 4C). These data suggest that fibroblasts from young individuals are under a basal level of oxidative stress in the *in vivo* oxygen environment that does not affect chromosomal stability. The relationship between oxidative stress and CIN was also examined in live cell imaging in the presence or absence of NAC. We confirmed that cells with chromosome missegregation, both lagging chromosomes and chromosome bridges, were reduced in fibroblasts from young and aged individuals in the presence of NAC (Fig. S2A, B). The duration of prometaphase and mitosis were reduced when cells were treated with NAC in fibroblasts from young and aged individuals (Fig. S2A, C), reflecting improved mitotic progression.

**Fig. 4.**
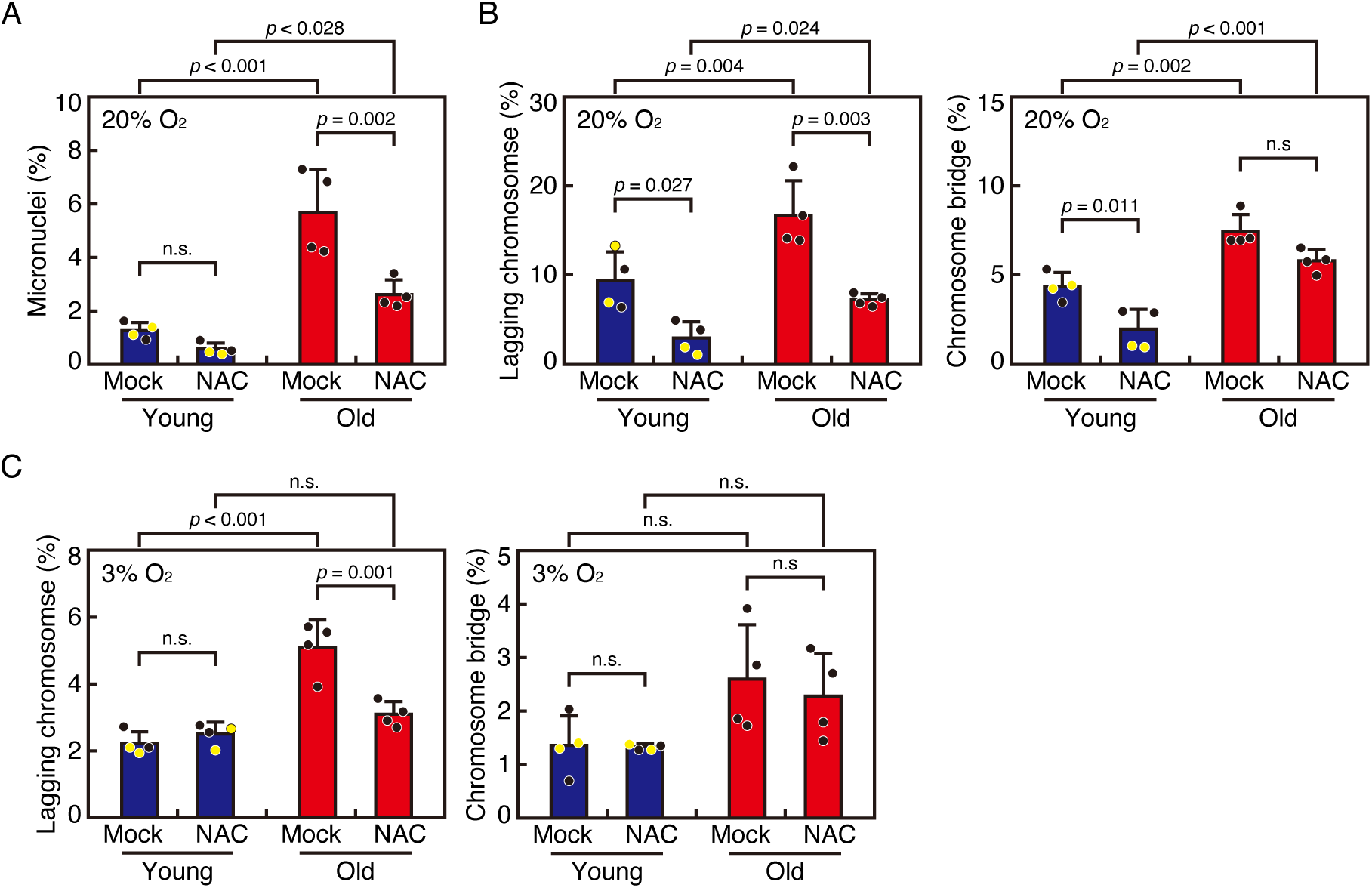
Antioxidant treatment ameliorates CIN in human fibroblasts. (A) Micronucleation rates of fibroblasts isolated from young and aged individuals in the presence or absence of an antioxidant. Fibroblasts from young and aged individuals (n = 4 each) were treated with NAC for 48 h. Then cells were fixed and stained with DAPI. Rates of micronucleation were quantified for 469-788 cells per cell line. Black-filled circles represent skin fibroblasts, while yellow-filled circles represent lung fibroblasts. Error bars represent S.D. P values were obtained using the Tukey–Kramer multiple comparison test. n.s., not statistically significant. (B) Chromosome missegregation rates of fibroblasts isolated from young and aged individuals in the presence or absence of NAC. Fibroblasts from young and aged individuals (n = 4 each) were treated as in (A). Rates of lagging chromosomes and chromosome bridges were quantified for 72-103 cells per cell line. Black-filled circles represent skin fibroblasts, while yellow-filled circles represent lung fibroblasts. Error bars represent S.D. P values were obtained using the Tukey–Kramer multiple comparison test. n.s., not statistically significant. (C) Chromosome missegregation rates of fibroblasts isolated from young and aged individuals cultured under 3% O_2_ in the presence or absence of NAC. Fibroblasts from young and aged individuals (n = 4 each) were cultured under 3% O_2_ and treated as in (A). Rates of lagging chromosomes and chromosome bridges were quantified for 51-157 cells per cell line. Black-filled circles represent skin fibroblasts, while yellow-filled circles represent lung fibroblasts. Error bars represent S.D. P values were obtained using the Tukey–Kramer multiple comparison test. n.s., not statistically significant.

### Antioxidant treatment ameliorates replication stress that is upstream of CIN in human fibroblasts

We next investigated the relationship between oxidative stress and replication stress in human fibroblasts. The number of γ-H2AX foci was reduced in fibroblasts from young and aged individuals after NAC treatment (Fig. 5A), which is consistent with previous reports that oxidative stress increases γ-H2AX foci (Katsube et al., 2014; Ye et al., 2016). We then observed 53BP1 foci and found that NAC treatment reduced the percentage of cells with 53BP1 nuclear bodies (Fig. 5B). Moreover, NAC treatment reduced the percentage of anaphase cells with UFBs (Fig. 5C), confirming that oxidative stress induces replication stress in human fibroblasts. NAC treatment reduced replication stress not only in fibroblasts from aged individuals, but also in those from young individuals, probably due to oxidative stress caused by the atmospheric oxygen environment, as shown in the previous section.

**Fig. 5.**
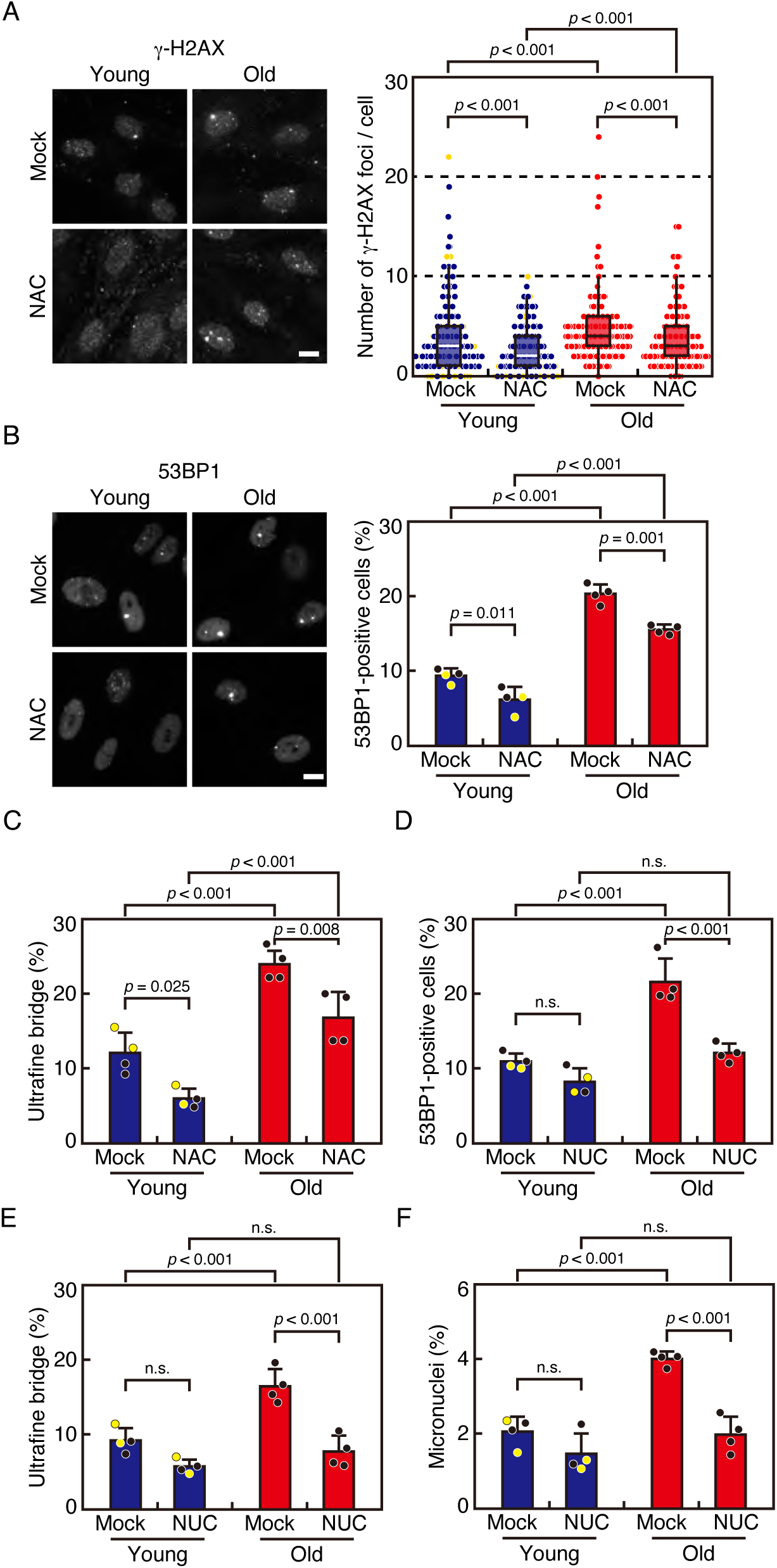
Antioxidant treatment ameliorates replication stress that is upstream of CIN in human fibroblasts. (A) Number of γ-H2AX foci in fibroblasts isolated from young and aged individuals in the presence or absence of NAC. Fibroblasts from young and aged individuals (n = 4 each) were treated with NAC for 48 h. Then cells were fixed and stained with an antibody against γ-H2AX. Representative images are shown in the left panels. Scale bar: 5 μm. Number of γ-H2AX foci were quantified for 33-49 cells per cell line and displayed as box and dot plots in the right graph. Red- and blue-filled circles represent skin fibroblasts, while yellow-filled circles represent lung fibroblasts. Error bars represent S.D. P values were obtained using the Steel-Dwass multiple comparison test. (B) Rates of cells with 53BP1 nuclear bodies in fibroblasts isolated from young and aged individuals in the presence or absence of NAC. Fibroblasts from young and aged individuals (n = 4 each) were treated with NAC for 48 h. Then cells were fixed and stained with an antibody against 53BP1. Cells with 53BP1 nuclear bodies were quantified for 526-770 cells per cell line. Black-filled circles represent skin fibroblasts, while yellow-filled circles represent lung fibroblasts. Error bars represent S.D. P values were obtained using the Tukey–Kramer multiple comparison test. (C) Rates of cells with ultrafine bridges in fibroblasts isolated from young and aged individuals in the presence or absence of NAC. Fibroblasts from young and aged individuals (n = 4 each) were treated with NAC for 48 h and stained with an antibody against PICH (ERCC6L). DNA was stained with DAPI. 72-103 cells were counted per cell line. Error bars represent S.D. Black-filled circles represent skin fibroblasts, while yellow-filled circles represent lung fibroblasts. P-value was obtained using the Tukey– Kramer multiple comparison test. (D) Rates of cells with 53BP1 nuclear bodies in fibroblasts isolated from young and aged individuals supplemented with or without nucleosides (Nuc). Fibroblasts from young and aged individuals (n = 4 each) were supplemented with nucleosides for 48 h. Then cells were fixed and stained with an antibody against 53BP1. 401-890 cells were counted per cell line. Error bars represent S.D. Black-filled circles represent skin fibroblasts, while yellow-filled circles represent lung fibroblasts. P values were obtained using the Tukey– Kramer multiple comparison test. n.s., not statistically significant. (E) Rates of cells with ultrafine bridges in fibroblasts isolated from young and aged individuals supplemented with or without nucleosides. Fibroblasts from young and aged individuals (n = 4 each) were treated as in (D). Then cells were fixed and stained with an antibody against PICH (ERCC6L). DNA was stained with DAPI. 39-70 cells were counted per cell line. Error bars represent S.D. Black-filled circles represent skin fibroblasts, while yellow-filled circles represent lung fibroblasts. P-value was obtained using the Tukey–Kramer multiple comparison test. n.s., not statistically significant. (F) Micronucleation rates of fibroblasts isolated from young and aged individuals supplemented with or without nucleosides. Fibroblasts from young and aged individuals (n = 4 each) were treated as in (D). Then cells were fixed and stained with DAPI. 521-741 cells were counted per cell line. Error bars represent S.D. Black-filled circles represent skin fibroblasts, while yellow-filled circles represent lung fibroblasts. P values were obtained using the Tukey–Kramer multiple comparison test. n.s., not statistically significant.

We then addressed the relationship between replication stress and CIN. It was reported that CIN is induced not only by defects in the mitotic process, but also by DNA damage in interphase caused by replication stress (Burrell et al., 2013; DePinho & Polyak, 2004), and we previously reported that replication stress causes CIN in fibroblasts from aged mice (Chen et al., 2023). To validate the causal relationship between replication stress and CIN in human fibroblasts, we supplemented the cells with nucleosides, which partially rescue replication stress (Burrell et al., 2013). In fibroblasts from aged individuals, the percentages of cells with 53BP1 nuclear bodies as well as UFBs were reduced (Fig. 5D, E), confirming that replication stress was alleviated by nucleoside treatment. Under this condition, the percentage of cells containing micronuclei was reduced in fibroblasts from aged individuals (Fig. 5F), suggesting that replication stress is a cause of CIN in these cells.

### Microtubules stabilization is a cause of CIN in fibroblasts from aged individuals

For proper chromosome segregation, kinetochores on the sister chromatids must attach to microtubules from opposite spindle poles, referred to as bi-orientation (Tanaka, 2012, 2013). Correction of erroneous kinetochore-microtubule attachments is crucial to ensure bi-orientation establishment for all chromosomes (Tanaka & Hirota, 2009). It was recently reported that replication stress causes CIN through stabilization of microtubules, which reduces the correction efficiency of erroneous kinetochore-microtubule attachments (Wilhelm et al., 2019). We previously reported that microtubules are stabilized in fibroblasts from aged mice, suggesting a link between microtubule stabilization and CIN (Chen et al., 2023). To examine the stability of microtubules in human fibroblasts, we quantified the intensity of spindle microtubules resistant to a 10-min cold treatment on ice, which depolymerizes unstable microtubules (Iemura & Tanaka, 2015). The intensity of spindle microtubules was comparable between fibroblasts from young and aged individuals at room temperature (Fig. 6A). However, spindle microtubule intensity after a 10-min cold treatment was significantly higher in fibroblasts from aged individuals (Fig. 6A), suggesting that microtubules were stabilized in these cells. The increase in the intensity of spindle microtubules was alleviated in the presence of NAC, indicating that oxidative stress is responsible for the microtubule stabilization (Fig. 6B, Fig. S3A). We then addressed whether increasing microtubule dynamics ameliorates CIN. To achieve this, we treated cells with UMK57, a potentiator of MCAK, a motor protein with microtubule depolymerizing activity (Orr et al., 2016). As shown in Fig. 6C and Fig. S3B, the intensity of spindle microtubules after a 10-min cold treatment was reduced by UMK57 treatment in cells from aged individuals, as expected. Under this condition, the percentage of cells containing micronuclei was reduced in cells from aged individuals, corroborating that microtubule stabilization is a cause of CIN (Fig. 6D). In contrast, the number of γ-H2AX foci as well as the percentage of cells containing 53BP1 nuclear bodies did not differ in the presence or absence of UMK57 (Fig. 6E, F), showing that replication stress is not downstream of microtubule stabilization. Collectively, these data suggest that oxidative stress causes microtubule stabilization, which is a cause of CIN in human fibroblasts with age.

**Fig. 6.**
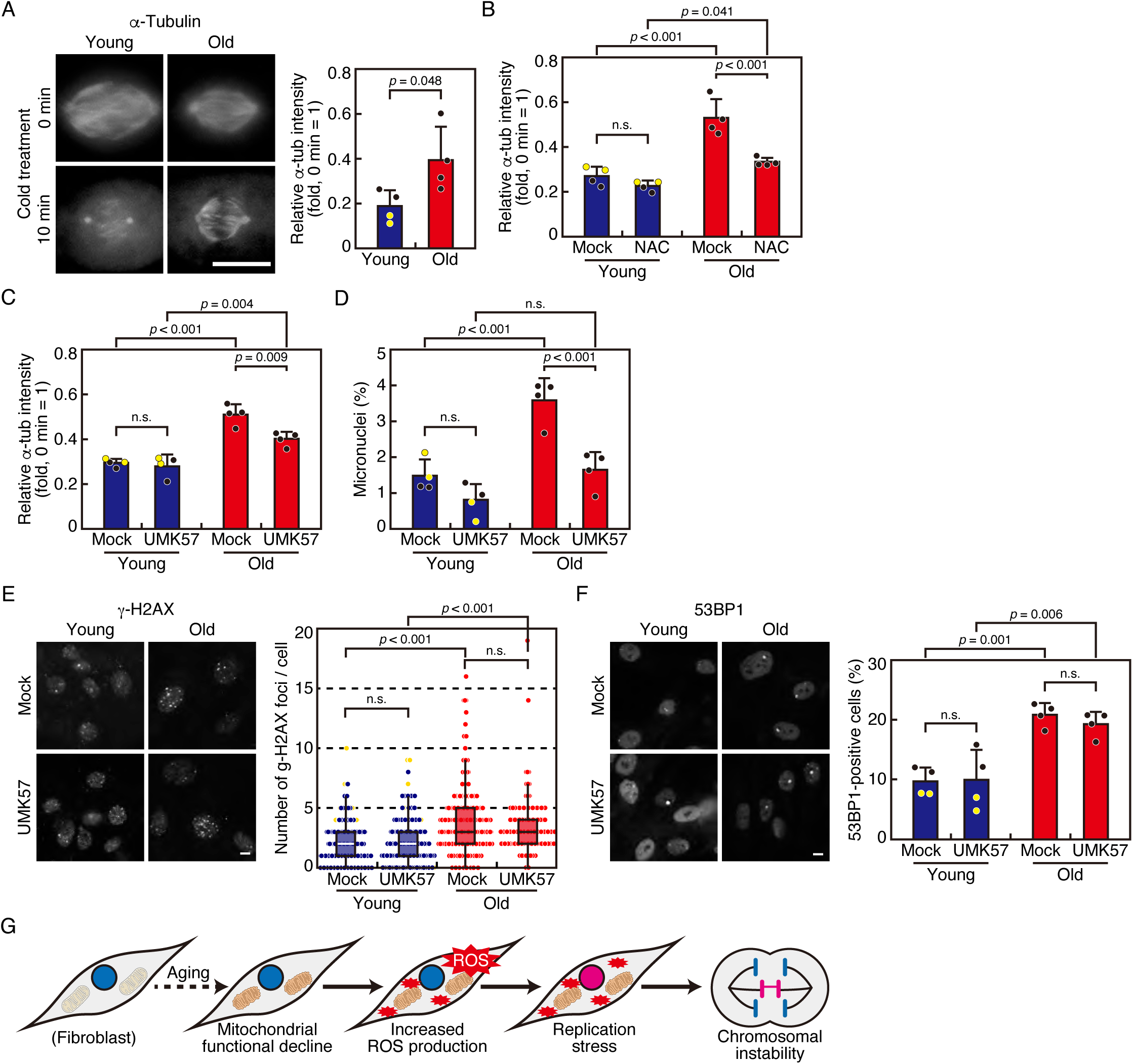
Microtubules stabilization is a cause of CIN in fibroblasts from aged individuals. (A) Microtubule stability in fibroblasts isolated from young and aged individuals. Fibroblasts from young and aged individuals (n = 4 each) were synchronized by RO-3306 treatment for 12 h, released and treated with MG132 for 3 h, and either directly fixed or fixed after cold treatment on ice for 10 min. Cells were stained with an antibody against α-tubulin, and relative fluorescence intensity on the spindle at 10 min compared with the intensity at 0 min was quantified for 5-7 cells per cell line. Representative images are shown in the left. Scale bar: 5 μm. Error bars in the right graph represent S.D. Black-filled circles represent skin fibroblasts, while yellow-filled circles represent lung fibroblasts. P-value was obtained using the Student’s *t*-test. (B) Microtubule stability in fibroblasts isolated from young and aged individuals in the presence or absence of NAC. Fibroblasts from young and aged individuals (n = 4 each) were treated with NAC for 48 h and treated as in (A). Relative fluorescence intensity on the spindle at 10 min compared with the intensity at 0 min was quantified for 10-24 cells per cell line. Error bars in the right graph represent S.D. Black-filled circles represent skin fibroblasts, while yellow-filled circles represent lung fibroblasts. P-value was obtained using the Tukey–Kramer multiple comparison test. n.s., not statistically significant. (C) Microtubule stability in fibroblasts isolated from young and aged individuals in the presence or absence of UMK57, an MCAK potentiator. Fibroblasts from young and aged individuals (n = 4 each) were treated with UMK57 for 24 h and treated as in (A). Relative fluorescence intensity on the spindle at 10 min compared with the intensity at 0 min was quantified for 10-28 cells per cell line. Error bars in the right graph represent S.D. Black-filled circles represent skin fibroblasts, while yellow-filled circles represent lung fibroblasts. P-value was obtained using the Tukey–Kramer multiple comparison test. n.s., not statistically significant. (D) Micronucleation rates of fibroblasts isolated from young and aged individuals in the presence or absence of UMK57. Fibroblasts from young and aged individuals (n = 4 each) were treated with UMK57 for 24 h. Then cells were fixed and stained with DAPI. 251-534 cells were counted per cell line. Black-filled circles represent skin fibroblasts, while yellow-filled circles represent lung fibroblasts. Error bars represent S.D. P values were obtained using the Tukey–Kramer multiple comparison test. n.s., not statistically significant. (E) Number of γ-H2AX foci in fibroblasts isolated from young and aged individuals in the presence or absence of UMK57. Fibroblasts from young and aged individuals (n = 4 each) were treated as in (D), and fixed and stained with an antibody against γ-H2AX. Representative images are shown in the left panels. Scale bar: 5 μm. Number of γ-H2AX foci were quantified for 49-70 cells per cell line and displayed as box and dot plots in the right graph. Red- and blue-filled circles represent skin fibroblasts, while yellow-filled circles represent lung fibroblasts. Error bars represent S.D. P values were obtained using the Steel-Dwass multiple comparison test. n.s., not statistically significant. (F) Cells with 53BP1 nuclear bodies in fibroblasts isolated from young and aged individuals in the presence or absence of UMK57. Fibroblasts from young and aged individuals (n = 4 each) were treated as in (D), and fixed and stained with an antibody against 53BP1. Representative images of 53BP1 nuclear bodies (arrowheads) are shown in the left panels. Scale bar: 5 μm. Cells with 53BP1 nuclear bodies were quantified for 261-416 cells per cell line. Black-filled circles represent skin fibroblasts, while yellow-filled circles represent lung fibroblasts. Error bars represent S.D. P values were obtained using the Tukey–Kramer multiple comparison test. n.s., not statistically significant. (G) Schematic of the mechanism of CIN in fibroblasts from aged individuals suggested in this study. See text for details.

## DISCUSSION

In the present study, we found that aneuploid cells increase with age in human fibroblasts. This is due to an increase in chromosome missegregation, namely CIN. Our analysis revealed that oxidative stress caused by increased ROS due to mitochondrial dysfunction is responsible for CIN with age. It was suggested that oxidative stress induces replication stress, which may cause CIN via microtubule stabilization (Fig. 6G). This is similar to the mechanism we previously demonstrated in primary mouse fibroblasts, suggesting that aging-related CIN induced by oxidative stress is a common phenomenon in mammalian cells. The presence of replication stress in fibroblasts from aged individuals was demonstrated not only by established hallmarks (53BP1 nuclear bodies and USBs), but also by direct measurement of replication fork progression using a DNA fiber assay, which was not shown in our previous study on cells from aged mice (Chen et al., 2023), further strengthening our conclusion.

In our previous study, we hypothesized that oxidative stress is a cause of CIN in primary mouse fibroblasts, since these cells immediately stopped proliferating under atmospheric oxygen conditions and exhibited an increase in chromosome missegregation (Chen et al., 2023). High-oxygen culture conditions lead to increased ROS production as a byproduct of the electron transport chain in mitochondria (Baez & Shiloach, 2014). In contrast, human fibroblasts can proliferate under atmospheric oxygen conditions and are considered resistant to oxidative stress (Hornsby, 2003). This difference is thought to be related to differences in the response to DNA damage, although the details are poorly understood (Hornsby, 2003). Despite the differences in the sensitivity to oxidative stress, our results show that the frequency of chromosome missegregation increases in human fibroblasts under atmospheric oxygen conditions, indicating that the appearance of CIN caused by oxidative stress is a common phenomenon even in human cells that are tolerant to oxidative stress. Mitotic defects caused by oxidative stress were also observed in human cells exposed to hydrogen peroxide (D’Angiolella et al., 2007; G. F. Wang et al., 2017). It has been reported that lowering the oxygen concentration promotes cell proliferation and mitotic progression in human cells (Kakudo et al., 2015; Stamatov et al., 2025). These findings suggest that the level of CIN observed in cells cultured under normal atmospheric oxygen conditions is overestimated compared to that in cells under physiological conditions in tissues. We confirmed that the chromosome missegregation rate in human fibroblasts was reduced under 3% oxygen concentration. However, fibroblasts from aged individuals show a higher rate of chromosome missegregation even under a 3% oxygen environment, which is consistent with previous reports of an increase in aneuploid cells with age (Ricke & van Deursen, 2013), and may suggest a causal relationship between oxidative stress and CIN with age in human cells.

NAC treatment improved all the phenotypes observed in fibroblasts from aged individuals we found in this study, including ROS levels, replication stress, and CIN, indicating that oxidative stress causes these phenotypes. Intriguingly, NAC treatment even improved mitochondrial function, as evidenced by an increase in mitochondrial membrane potential, as reported previously (Zheng et al., 2024; Zhou et al., 2021), which would further contribute to reducing ROS levels. We also found that microtubule stabilization was ameliorated by NAC treatment, which was not shown in our previous study (Chen et al., 2023), confirming the causal relationship between oxidative stress and microtubule stabilization. However, NAC treatment did not improve the parameters in fibroblasts from aged cells to the level comparable to those in fibroblasts from young individuals, suggesting that oxidative stress does not explain all the changes seen in fibroblasts from aged individuals.

As a cause of CIN, we found that microtubules are stabilized in fibroblasts from aged individuals, which may be downstream of replication stress, as reported previously. (Wilhelm et al., 2019). Microtubule stabilization in fibroblasts from elderly people and restoration of CIN by increasing microtubule dynamics was also observed in a previous study (Barroso-Vilares et al., 2020). According to a report from the same group (Macedo et al., 2018), a correlation exists between increased chromosome missegregation in fibroblasts of aged individuals and the downregulation of FoxM1, a transcription factor that plays a role in mitotic gene expression. The relationship between FoxM1 repression and oxidative stress in the development of CIN requires further investigation.

What are the consequences of the age-related CIN? Oxidative stress and the DNA damage it causes are known to induce cellular senescence (d’Adda di Fagagna, 2008), which is one of the hallmarks of aging and related to aging-related disorders (Lopez-Otin et al., 2013). It has also been suggested that aneuploidization, resulting from mitotic defects in cells from elderly people, triggers the full senescence phenotype (Macedo et al., 2018). CIN is also known not only as a consequence of cancer, but also as a cause (Tanaka & Hirota, 2016). Because oxidative stress is a known cause of CIN in cancer (Kudryavtseva et al., 2016), the link we discovered between oxidative stress and CIN in cells from aged individuals could be connected to age-related oncogenesis. The higher sensitivity of mouse cells to oxidative stress compared to human cells may be related to their higher susceptibility to oncogenic transformation. Interestingly, when mouse embryonic fibroblasts with oncogene-induced CIN were treated with antioxidants, the CIN was reduced, implying a role of oxidative stress in CIN induction during neoplastic transformation (Woo & Poon, 2004). Our finding that the modal number of chromosomes did not change in fibroblasts from aged individuals suggests that the aneuploid cells resulting from chromosome missegregation do not propagate to alter the karyotype of the cell population, supposedly due to the mechanism to prevent the proliferation of aneuploid cells, in which p53 plays a pivotal role (Thompson & Compton, 2010). When p53 is lost or inactivated, the age-related CIN may lead to oncogenesis by propagating aneuploid cells.

The finding that oxidative stress induces CIN in human cells as well as in mice during aging through a similar mechanism is important for understanding the background of aging and oncogenesis in humans. As a certain level of ROS is necessary for cellular homeostasis (Lopez-Otin et al., 2013), further study is required to regulate ROS level within a proper range in order to maintain chromosomal stability and prevent age-related disorders including cancer.

## MATERIALS AND METHODS

### Cell culture and synchronization

The following human fibroblasts were obtained from the NIA Aging Cell Repository in the Coriell Institute for Medical Research (https://www.coriell.org): GM03349 (skin fibroblast, 10yo), GM08398 (skin fibroblast, 8yo), AG04434 (lung fibroblast, 19fw), AG06556 (lung fibroblast, 20fw), AG10884 (skin fibroblast, 87yo), AG11017 (skin fibroblast, 87yo), AG13129A (skin fibroblast, 89yo), and AG05248D (skin fibroblast, 87yo). Cells were cultured in Eagle’s Minimum Essential Medium with Earle’s salts and non-essential amino acids with 2mM L-glutamine supplemented with 15% fetal bovine serum (FBS) at 37℃ in 5% CO_2_. Population doubling level (PDL) was calculated using the formula PDL = 3.33 × [log (number of cells harvested) – log (number of cells seeded)] + (doubling level at seeding). Cells at PDL (population doubling level) ranging from 17 to 35, during which the cells grew at a constant rate, were used for experiments. To enrich mitotic cells, cells were treated with 2 mM of thymidine (Wako) for 24 h, followed by treatment with 10 μM of RO-3306 (Tokyo Chemical Industry, R0201) for 12 h, then released for indicated periods before fixation.

### Reagents and antibodies

N-acetyl-L-cysteine (NAC; Merck, A7250) was used at 5 mM. Nocodazole (Merck, M1404) and Eg5 inhibitor III (Merck, 324622) were used at 1 μM. MG-132 (Merck, M8699) was used at 20 μM. UMK57 (AOBIOS, AOB8668) was used at 1 μM. For nucleoside supplementation, a mix of deoxycytidine (Tokyo Chemical Industry, D3583), deoxyadenosine (Tokyo Chemical Industry, D0046), thymidine (Tokyo Chemical Industry, T0233) and deoxyguanosine (Tokyo Chemical Industry, D0052) was applied at 20 μM for 48 h. A rabbit polyclonal antibody for 53BP1 (NOVUS Biologicals, NB100-304) and PICH (ERCC6L; Proteintech, 15688-1-AP) were used for immunofluorescence at 1:1,000. A mouse monoclonal antibody for α-tubulin (B5-1-2; Merck, T5168), alpha smooth muscle actin (α-SMA, 1A4; Abcam, ab7817), and phospho-Histone H2A.X (Ser139 (γ-H2AX), JBW301; Merck, 05-636) were used for immunofluorescence at 1:1,000.

### Metaphase chromosome spreads

Cells were treated with 2 mM of thymidine for 24 h, followed by treatment with 10 μM of RO-3306 for 12 h, then treated with 20 μM of MG-132 and 2 μM of nocodazole for 2 h. Mitotic cells collected by a shake-off were swollen hypotonically by adding 4 volumes of tap water for 5 min, and then fixed with Carnoy’s solution (methanol: acetic acid = 3:1). The cell suspension was spotted onto slide glasses, dried, and mounted with VECTASHIELD Mounting Medium with DAPI (Vector Laboratories, H-1200). Images were captured on an Olympus IX-83 inverted microscope (Olympus) controlled by cellSens Dimension 1.18 (Olympus) using a ×100 1.40 numerical aperture (NA) UPlanS Apochromat oil objective lens (Olympus).

### Immunofluorescence analysis

Immunofluorescence analysis was performed according to a previous report (Iemura & Tanaka, 2015). Briefly, cells were grown on a glass coverslip and fixed with 3% paraformaldehyde in PBS for 15 min at room temperature, and permeabilized with 1% Triton X-100 in PBS for 5 min. Fixed cells were incubated with primary antibodies for more than 12 h at 4℃, washed three times with PBS supplemented with 0.02% Triton X-100, and incubated with secondary antibodies coupled with Alexa-Fluor-488/594 (Thermo Fisher Scientific, 1:2,000) for 1 h at room temperature. Antibody incubations were performed in PBS supplemented with 0.02% Triton X-100. After final washes, cells were mounted with VECTASHIELD Mounting Medium with DAPI. Z-image stacks were captured in 0.2 or 0.27 μm increments on an Olympus IX-83 inverted microscope controlled by cellSens Dimension 1.18 using a ×100 1.40 or ×60 1.35 NA UPlanS Apochromat oil objective lens. Deconvolution was performed when necessary. Image stacks were projected and saved as TIFF files. All cells analyzed were selected from non-overlapping fields.

### SPiDER-βGal staining

To observe the presence of micronuclei together with SA-β-gal activity, cells were treated with 1 μM SPiDER-βGal (Dojindo, SG02) for 15 min, and fixed with 3% paraformaldehyde in PBS for 15 min at room temperature. Then cells were stained with DAPI for 20 min. Images were captured on an Olympus IX-83 inverted microscope controlled by cellSens Dimension 1.18 using×60 1.35 NA Plan Apochromat oil objective lens.

### DNA fiber assay

DNA fiber assay was performed as reported previously (Luebben et al., 2014). Briefly, cells were incorporated with 200 μM digoxigenin-dUTPs (Roche) for 10 min, followed by a gentle wash with fresh pre-warmed medium and the second incorporation of 200μM biotin-dUTPs (Roche) for 10 min. After labeling, cells were trypsinized and fixed (acetic acid:methanol = 1:3). Then the fixed cell suspension is dropped onto microscope slides and air-dried. To extend the DNA fibers, the slides are immersed in lysis buffer (0.5 % SDS, 50 mM EDTA, 200 mM Tris–HCl, pH 7.0), then placed in a high-humidity chamber. After extension, the slides are washed, fixed, and dried. For detection, slides are treated with 1 x blocking reagent (Roche), followed by the addition of detection buffer containing Anti-Digoxigenin-Rhodamine, Fab fragments from sheep: (Roche) and Streptavidin, Alexa Fluor^®^ 488 Conjugate (Life technologies) to visualize the labeled DNA forks. The slides are then washed and mounted with DAPI-supplemented medium, making them ready for microscopic analysis.

### ROS detection

Cells were grown in 3.5cm glass bottom dishes (MatTek), and ROS was measured using CellROX Oxidative Stress Reagents (Molecular Probes, C10444), according to the manufacturer’s instructions. Briefly, the cells were stained with 5 μM CellROX green reagents with 50 μM verapamil hydrochloride (Fujifilm Wako Chemicals, 222-00781) for 30 min at 37℃ in 5% CO_2_ before imaging. Superoxide in mitochondria was detected using MitoSOX Red (Molecular Probes, M36008), according to the manufacturer’s instructions. Briefly, the cells were stained with 1 μM MitoSOX Red with 50 μM verapamil hydrochloride for 10 min at 37℃ in 5% CO_2_ before imaging. Images were captured on an Olympus IX-83 inverted microscope controlled by cellSens Dimension 1.18 using ×60 1.40 NA Plan Apochromat oil objective lens. Image stacks were projected and saved as TIFF files.

### Mitochondrial membrane potential assay

Mitochondrial membrane potential was measured using JC-10 (AAT Bioquest, 22204) according to the manufacturer’s instructions. Briefly, the cells were grown in 3.5cm glass bottom dishes and stained with 1× JC-10 with 50 μM verapamil hydrochloride for 30 min at 37℃ in 5% CO_2_ before imaging. Images were captured on an Olympus IX-83 inverted microscope controlled by cellSens Dimension 1.18 using ×60 1.40 NA Plan Apochromat oil objective lens. Image stacks were projected and saved as TIFF files.

### Live cell imaging

Cells were grown in 12 or 6-well plates (Violamo) and treated with 100 nM SiR-DNA (Cytoskeleton, CY-SC007) and 2 μM verapamil hydrochloride for 4 h at 37℃ in 5% CO_2_ before imaging. Recordings were made every 2.5 min for 48 h using Celldiscoverer 7 live cell imaging system (Carl Zeiss) controlled by ZEN 2.6 (Carl Zeiss) at 37°C in 5% CO_2_ and 20% O_2_ using ×20 0.7 NA Plan Apochromat objective lens (Carl Zeiss). For observation of cells after mitotic arrest, cells treated with 1 μM nocodazole or Eg5 inhibitor III were observed every 15 min for 96 h. Images were analyzed using ZEN 2.6.

### Statistical analysis

The Mann–Whitney *U*-test was used for comparison of dispersion, and a two-sided *t*-test was used for comparisons of average. A two-sided F-test validated the dispersibility of each category before the Student’s *t*-test. If the result of the F-test was an unequal variance, a significant difference between samples was validated by a two-sided Welch’s *t*-test. A one-way ANOVA test was used with the Tukey–Kramer post hoc test for comparisons between all groups showing normal distribution. The Kolmogorov–Smirnov test verified the normality of data distribution for each group before the one-way ANOVA test. If the result of the Kolmogorov–Smirnov test was a non-nominal distribution, the significant differences between all groups were validated by the Kruskal–Wallis test, which was used with Steel–Dwass post hoc test. A Chi-squared test was used for comparison between the measured value and theoretical value. All statistical analyses were performed with EZR (Kanda, 2013), which is a graphical user interface for R (R Core Team, R: A language and environment for statistical computing, https://www.R-project.org/, (2018)). More precisely, it is a modified version of R commander designed to add statistical functions frequently used in biostatistics. Samples for analysis in each dataset were acquired in the same experiment, and all samples were calculated at the same time for each dataset.

## Acknowledgements

The authors thank members of the K.T. laboratory for discussions, and H. Sato for technical assistance.

## Author contributions

K.Z. and K.T. designed the experiments. K.Z., G.C., and R.I. performed the experiments. K.Z., K.I., and K.T. wrote the manuscript. K.T. supervised the work.

---

**Fig. S1.**
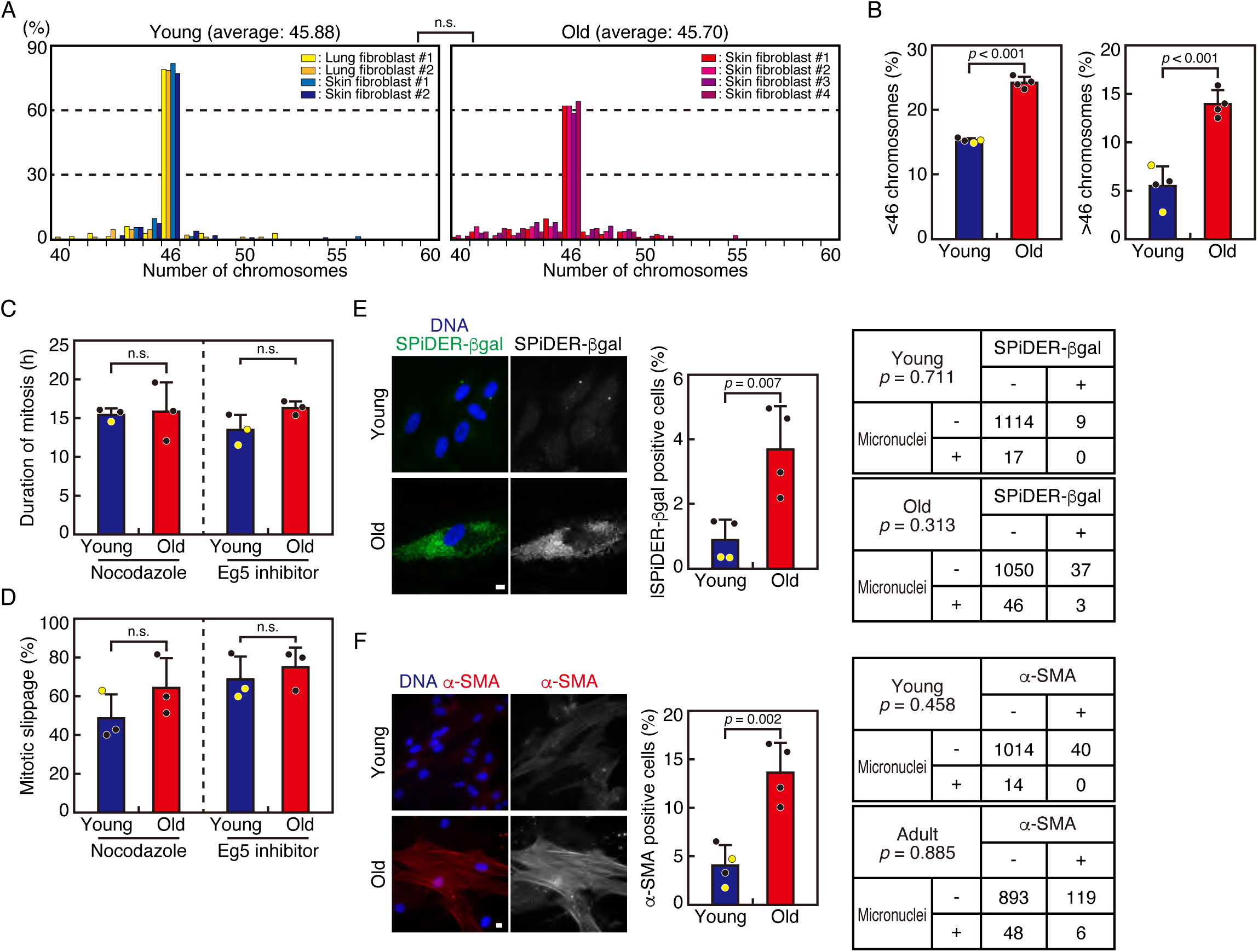
Fibroblasts isolated from aged human exhibit CIN. (A) Number of chromosomes in skin fibroblasts isolated from young and aged individuals. Number of chromosomes in 50-105 metaphase chromosome spreads of fibroblasts isolated from young (left) and aged (right) individuals (n = 4 each) are shown. Error bars represent S.D. P values were obtained using the Steel–Dwass multiple comparison test. n.s., not statistically significant. (B) Percentages of cells with fewer (left) and more than (right) 46 chromosomes isolated from young and aged individuals shown in (A). Black-filled circles represent skin fibroblasts, while yellow-filled circles represent lung fibroblasts. Error bars represent S.D. P-values were obtained using the Student’s *t*-test. (C) Duration of mitosis in fibroblasts isolated from young and aged individuals in the presence of nocodazole or an Eg5 inhibitor. Fibroblasts from young and aged individuals (n = 3 each) were treated with 1 μM nocodazole or 1 μM Eg5 inhibitor III. Cells were observed for 96 h and the duration of mitosis, which were determined as the duration from mitotic cell rounding to cell rupture (mitotic cell death) or cell flattening (mitotic slippage), was quantified for 31-71 cells per cell line. Black-filled circles represent skin fibroblasts, while yellow-filled circles represent lung fibroblasts. Error bars represent S.D. P values were obtained using the Student’s *t*-test. n.s., not statistically significant. (D) Rate of mitotic slippage in fibroblasts isolated from young and aged individuals in the presence of nocodazole or an Eg5 inhibitor. Fibroblasts from young and aged individuals (n = 3 each) treated and observed as in (C) were categorized as either they underwent mitotic cell death or mitotic slippage, and the percentages of cells that underwent mitotic slippage are shown. Black-filled circles represent skin fibroblasts, while yellow-filled circles represent lung fibroblasts. Error bars represent S.D. P values were obtained using the Student’s *t*-test. n.s., not statistically significant. (E) Relationship between SPiDER-βGal-positive cells and cells with micronuclei in fibroblasts isolated from young and aged individuals. Fibroblasts from young and aged individuals (n = 4 each) were fixed and SPiDER-βGal signal and micronuclei were detected. Representative images are shown. Scale bar: 5 μm. Black-filled circles represent skin fibroblasts, while yellow-filled circles represent lung fibroblasts. Error bars represent S.D. P values were obtained using the Student’s *t*-test. Presence or absence of SPiDER-βGal signal and micronuclei were quantified for 273-302 cells per cell line and shown in 2×2 contingency tables in the right. P-values were obtained using the chi-square test. (F) Rates of α-SMA-positive cells in fibroblasts isolated from young and aged individuals. Fibroblasts from young and aged individuals (n = 4 each) were fixed and stained with an antibody against α-SMA. DNA was stained with DAPI. Representative images are shown. Scale bar: 5 μm. Rates of α-SMA-positive cells were quantified for 218-306 cells per cell line. Black-filled circles represent skin fibroblasts, while yellow-filled circles represent lung fibroblasts. Error bars represent S.D. P values were obtained using the Student’s *t*-test. Presence or absence of α-SMA signal and micronuclei were quantified for 218-306 cells per cell line and shown in 2×2 contingency tables. P-values were obtained using the chi-square test.

**Fig. S2.**
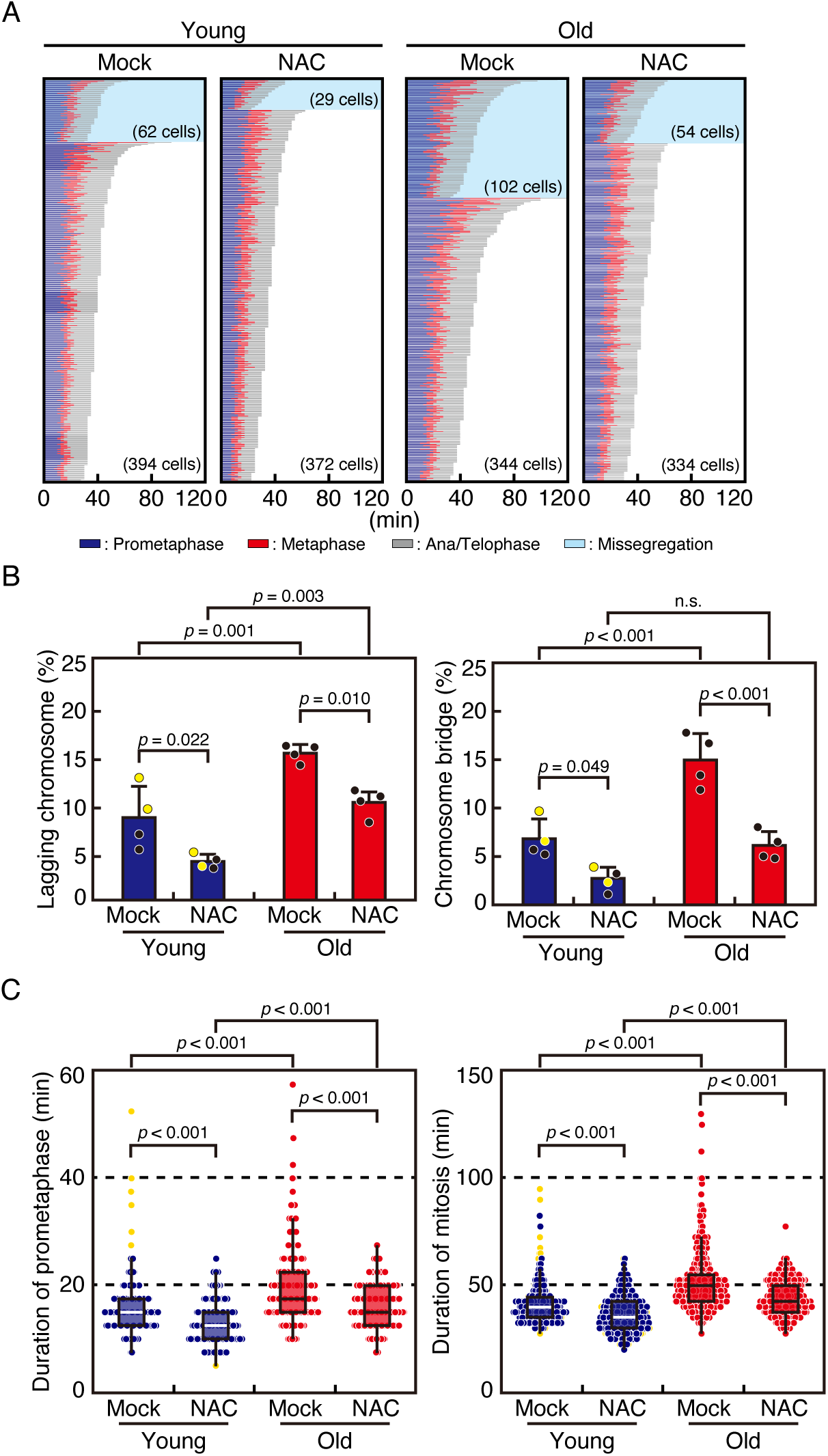
Antioxidant treatment ameliorates CIN in human fibroblasts. (A) Mitotic progression of fibroblasts isolated from young and aged individuals in the presence or absence of NAC. Fibroblasts from young and aged individuals (n = 4 each) were treated with SiR-DNA with or without NAC and subjected to live cell imaging for 96 h. Images were taken every 2.5 min, and mitotic progression of individual cells was tracked and displayed in different colors depending on the mitotic phases. Cells that underwent chromosome missegregation are separately shown, and number of cells in each category is indicated. (B) Chromosome missegregation rates of fibroblasts isolated from young and aged individuals in the presence or absence of NAC. Rates of lagging chromosomes and chromosome bridges in fibroblasts from young and aged individuals shown in (A) are plotted. Black-filled circles represent skin fibroblasts, while yellow-filled circles represent lung fibroblasts. Error bars represent S.D. P values were obtained using the Tukey–Kramer multiple comparison test. (C) Mitotic duration of fibroblasts isolated from young and aged individuals in the presence or absence of NAC. Time from nuclear envelope breakdown to cytokinesis (duration of prometaphase), and time from nuclear envelope breakdown to cytokinesis (duration of mitosis) in fibroblasts from young and aged individuals shown in (A) are plotted. Red- and blue-filled circles represent skin fibroblasts, while yellow-filled circles represent lung fibroblasts. Error bars represent S.D. P values were obtained using the Steel-Dwass multiple comparison test.

**Fig. S3.**
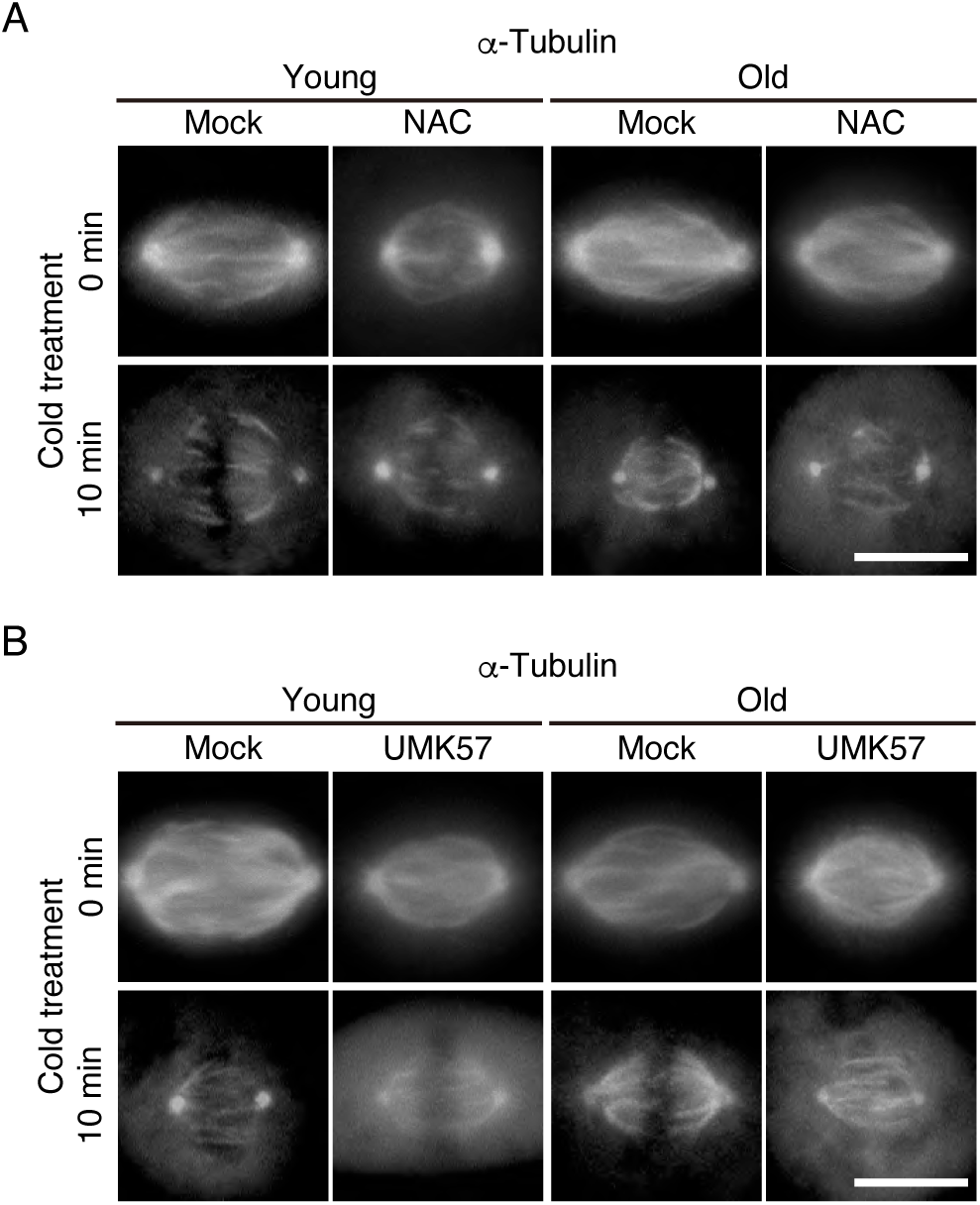
Microtubule stability in fibroblasts isolated from young and aged individuals in the presence or absence of NAC or UMK57. (A) Representative images of the spindles before or after 10 min cold treatment in fibroblasts treated with or without NAC. Cells were treated as in Fig. 5B. Scale bar: 5 μm. (B) Representative images of the spindles before or after 10 min cold treatment in fibroblasts treated with or without UMK57. Cells were treated as in Fig. 5C. Scale bar: 5 μm.

